# A set informative multiple autosomal markers for human identification: forensic research and population genetics analysis in a Chinese Xinjiang Hui group

**DOI:** 10.1101/197772

**Authors:** Tong Xie, Yuxin Guo, Ling Chen, Yating Fang, Yunchun Tai, Yongsong Zhou, Pingming Qiu, Bofeng Zhu

## Abstract

In recent years, insertion/deletion (InDel) markers became a promising and useful supporting tool in forensic identification cases and biogeographic research field. In this study, 30 InDel loci were explored to reveal the genetic diversities and genetic relationships between Chinese Xinjiang Hui group and the 24 previously studied populations using varies methods such as forensic statistical parameter analysis, phylogenetic reconstruction, STRUCTURE analysis, multi-dimensional scaling, and principal component analysis. The observed heterozygosity and expected heterozygosity ranged from 0.1971 (HLD118) to 0.5092 (HLD 92), 0.2222 (HLD 114) to 0.5000 (HLD 6), respectively. Besides, after Bonferroni correction, no deviations from Hardy-Weinberg equilibrium tests were found at all 30 loci in Xinjiang Hui group. The cumulative probability of exclusion and combined discrimination power were 0.988849 and 0.99999999999378, respectively, which indicated that the 30 loci could be used as complementary genetic markers for paternity test and be qualified for personal identification in forensic cases. In this study, we found that Xinjiang Hui group had close relationships with most Chinese groups, especially Han populations, and all the results based on different genetic methods we used had a strong support for this finding. The 30 InDel loci has important significance in forensic identification research, in spite of this, for a better understanding of genetic background of the Chinese Xinjiang Hui group, molecular genetic genotyping at various genetic markers is necessary in future studies.

**Summary Statement:** We report here, a promising Individual identification and population differentiation maker which could be used in forensic cases.

## 1 Introduction

In 2002, Weber et al. first reported human diallelic InDel polymorphisms (Weber et al., 2002) which made InDels become promising genetic markers in the forensic identification cases and biogeographic inference of biological samples. From then on, a large number of studies have been published on the basis of InDels including the identification of ancestry affiliation, reconstruction the genetic structure of different human populations (Pereira et al., 2009) and practical case applications in various forensic cases. In recent years, InDels were regarded as new tremendous potential diallelic genetic markers in population genetic studies and forensic sciences because those markers combined the advantages of both STRs and SNPs: extensive distribution, short amplicon size and low mutation rate (Rui et al., 2009, Fondevila et al., 2012). Besides, InDels also showed similar length polymorphisms similar with STR and which made it possible to be detected by the current forensic DNA testing platforms such as capillary electrophoresis. Because of the short amplification, InDels can be successfully used in highly degraded DNA samples. There is at least one InDel in average 7.2 kb in human genome according to the milestone research of Mills et al [5]. In addition, InDel comprises about 20% of all human DNA polymorphisms (RE et al., 2006) which makes it to be preferential select for forensic applications. In general, InDel played an important role as effective supplement of existing genetic markers in personal identification and biogeographic ancestry analyses.

According to the previous study, the Hui group is one of the most widespread ethnic groups in China [25]. The Hui group live in many areas in China such as Ningxia, Gansu, Qinghai, Xinjiang, Henan, Anhui, Liaoning, Heilongjiang, Shaanxi provinces, and so on. Besides, modern archaeological record [29] revealed that during the initial period of the 13th century, the Persians, Arabs and Islamic-oriented people made their migration to China due to voluntary or war. Because those people had the identical religious beliefs with people living in northwest area, so they were called Hui or Huihui and considered as the ancestors of contemporary Hui group [29]. This study choose Xinjiang Hui group as research subject, and used 30 InDel loci to evaluate the forensic parameters and population genetic differentiations between Xinjiang Hui group and other 24 populations.

## 2 Material and Methods

### 2.1 Sample collections and DNA extraction

We collected bloodstain specimen from 487 unrelated healthy Hui individuals living in Xinjiang province, China. In this study, donors that there were no blood relationships with each other should have unrelated ancestors living in Xinjiang province for over three generations and have no significant migration in their family history. The collection process was followed by the human and ethical research principles of Southern Medical University, China. Human genomic DNA was isolated from bloodstain samples utilizing the method of Chelex-100.

### 2.2 PCR amplification and InDels genotyping

In this study, we conducted the multiple PCR amplification with fluorescent of autosomal 30 InDels by one commercial kit, Qiagen Investigator DIPplex reagent (Qiagen, Hilden, Germany) and performed the PCR amplification in a GeneAmp PCR System 9700 Thermal Cycler (Applied Biosystems, Foster City, CA, USA) following the manufacturer’s instruction. The genotyping of 30 InDels were conducted by capillary electrophoresis on ABI 3500xl Genetic Analyzer (Applied Biosystems, Foster City, CA, USA) and analyzed by GeneMapper *ID –X* v 1.5 (Applied Biosystems, Foster City, CA, USA).

### 2.3 Statistical analysis

The linkage disequilibrium (LD) analysis was carried based on allele frequencies by the SNP Analyzer version 2.0 (Istech, South Korea). The forensic statistical parameter such as the expected heterozygosity (He), the observed heterozygosity (Ho), the polymorphic information content (PIC), power of discrimination (PD), power of exclusion (PE), typical paternity index (TPI) and Hardy-Weinberg equilibrium (HWE) were calculate by modified PowerStates (version 1.2) spreadsheet (Promega, Madison, WI, USA), and the two Heat maps were conducted by *R* statistical software v3.0.2. The analysis of molecular variance (AMOVA) was conducted by Arlequin (version 3.0). The multi-dimensional scaling (MDS) of the populations was conducted by SPSS software (version 18.0). In addition, on the basis of the 30 InDels allele frequencies, the DA distance was calculated by (DISPAN) program, what’s more the phylogenetic reconstruction was performed by MAGA software (version5.0) (based on DA distance) and PHYLIP software v 3.6, respectively (based on allele frequency), respectively, and. Principal component analysis (PCA) was carried by the software of EIGENSOFT 6.0.1 (based on DA distances of pairwise populations), while the population structure was performed by the STRUCTURE program (version 2.3.4).

## 3 Results

### 3.1 Linkage disequilibrium analysis of 30 InDels

To evaluate LD for all possible combination between the 30 InDel loci, we used the SNP Analyzer program. As shown in Supplementary Fig 1, the pairwise linkage disequilibrium was displayed in the all square grids with no crimson color covered by darker red curve existing in those grids, and the values of *r*^2^ less than 0.8 (data not shown). The result indicated that any two different InDels showed no strong linkage disequilibrium (Barrett).

**Fig 1.**
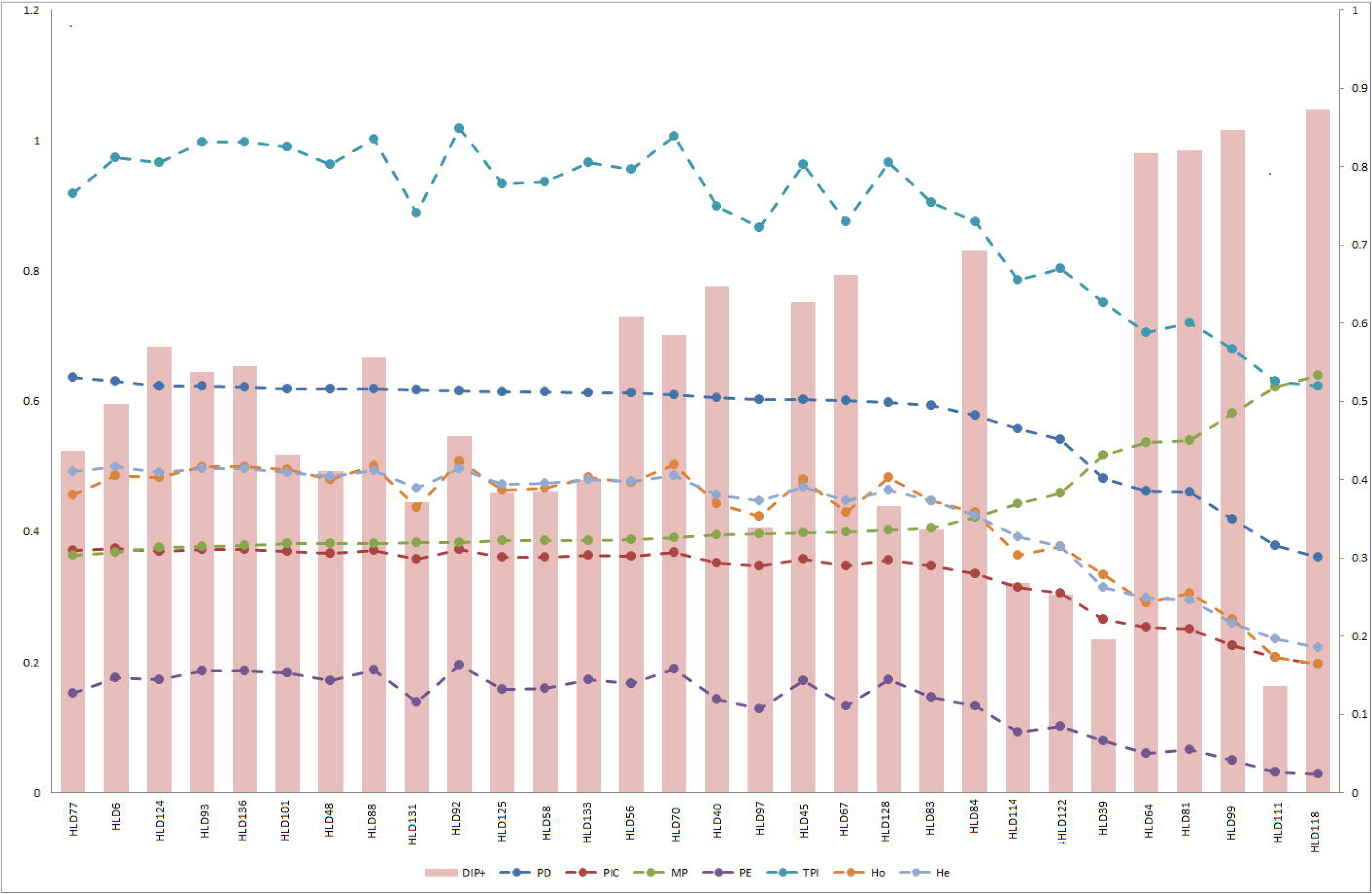
Plots of allele frequency and forensic parameters of the Chinese Xinjiang Hui group on account of 30 InDel loci.

### 3.2 Forensic statistical parameters of 30 InDels

Allele frequencies and the forensic efficiency parameters of 30 InDels in the Chinese Xinjiang Hui group were shown in Fig 1 and Table 1, respectively. In Fig 1, the Y-axis represented the forensic efficiency parameters and the right Y-axis represented allele frequencies. After applying Bonferroni correction (p=0.05/30) for all 30 InDel loci, the HWE tests were conducted and we obtained the *p* values ranging from 0.1042 (HLD 77) to 0.9957 (HLD 122), and the HWE tests showed no significant deviation from the expected value (*p*>0.0017).Among the 30 InDel loci, the deletion allele frequencies varied from 0.1273 (HLD 118) to 0.8634 (HLD 111), with a mean value of 0.4983. The Ho and He ranged from 0.1971 (HLD118) to 0.5092 (HLD 92), 0.2222 (HLD 114) to 0.5000 (HLD 6) with the mean value of 0.4239 and 0.4283, respectively. The MP, TPI and PIC values were in the range of 0.3637 (HLD 77) to 0.6390 (HLD 118); 0.6228 (HLD 118) to 1.0188 (HLD 92; and 0.1975 (HLD 118)to 0.3750(HLD 6), with the mean value of 0.4290, 0.8855 and 0.3330, respectively. The PD varied from 0.3610 (HLD 118) to 0.6363 (HLD 77), separately. The highest value of PE was 0.1957 at HLD 92 locus, while the lowest value was 0.0290 at HLD 118 locus, and the mean value is 0.1378. In this study, the lowest Ho, He, PIC, TPI, PD and PE values were observed at HLD 118 locus, and in the preceding studied groups, HLD 118 locus also showed the lowest polymorphisms (Guo et al., 2016).

**Table 1.** The allele frequencies andfor 30 InDel in Chinese Xinjiang Hui Group (n=487).

| HLD | DIP(-) | DIP(+) | PD | MP | PIC | PE | TPI | He | Ho | p |
| --- | --- | --- | --- | --- | --- | --- | --- | --- | --- | --- |
| HLD6 | 0.5041 | 0.4959 | 0.6314 | 0.3686 | 0.3750 | 0.1761 | 0.9740 | 0.5000 | 0.4867 | 0.5417 |
| HLD39 | 0.8039 | 0.1961 | 0.4820 | 0.5180 | 0.2656 | 0.0788 | 0.7515 | 0.3153 | 0.3347 | 0.3647 |
| HLD40 | 0.3532 | 0.6468 | 0.6053 | 0.3947 | 0.3525 | 0.1427 | 0.8985 | 0.4569 | 0.4435 | 0.5402 |
| HLD45 | 0.3737 | 0.6263 | 0.6023 | 0.3977 | 0.3585 | 0.1710 | 0.9625 | 0.4681 | 0.4805 | 0.5985 |
| HLD48 | 0.5893 | 0.4107 | 0.6182 | 0.3818 | 0.3669 | 0.1710 | 0.9625 | 0.4840 | 0.4805 | 0.8581 |
| HLD56 | 0.3922 | 0.6078 | 0.6127 | 0.3873 | 0.3631 | 0.1677 | 0.9549 | 0.4768 | 0.4764 | 0.9697 |
| HLD58 | 0.6150 | 0.3850 | 0.6138 | 0.3862 | 0.3614 | 0.1595 | 0.9365 | 0.4736 | 0.4661 | 0.7262 |
| HLD64 | 0.1828 | 0.8172 | 0.4628 | 0.5372 | 0.2541 | 0.0601 | 0.7058 | 0.2987 | 0.2916 | 0.7201 |
| HLD67 | 0.3378 | 0.6622 | 0.6003 | 0.3997 | 0.3473 | 0.1327 | 0.8759 | 0.4474 | 0.4292 | 0.4073 |
| HLD70 | 0.4158 | 0.5842 | 0.6093 | 0.3907 | 0.3678 | 0.1902 | 1.0062 | 0.4858 | 0.5031 | 0.4594 |
| HLD77 | 0.5626 | 0.4374 | 0.6363 | 0.3637 | 0.3710 | 0.1517 | 0.9189 | 0.4922 | 0.4559 | 0.1042 |
| HLD81 | 0.1797 | 0.8203 | 0.4603 | 0.5397 | 0.2513 | 0.0660 | 0.7204 | 0.2948 | 0.3060 | 0.5988 |
| HLD83 | 0.6632 | 0.3368 | 0.5938 | 0.4062 | 0.3469 | 0.1456 | 0.9052 | 0.4467 | 0.4476 | 0.9831 |
| HLD84 | 0.3070 | 0.6930 | 0.5784 | 0.4216 | 0.3350 | 0.1327 | 0.8759 | 0.4255 | 0.4292 | 0.8853 |
| HLD88 | 0.4435 | 0.5565 | 0.6181 | 0.3819 | 0.3718 | 0.1884 | 1.0021 | 0.4936 | 0.5010 | 0.7608 |
| HLD92 | 0.5441 | 0.4559 | 0.6164 | 0.3836 | 0.3730 | 0.1957 | 1.0188 | 0.4961 | 0.5092 | 0.5773 |
| HLD93 | 0.4630 | 0.5370 | 0.6228 | 0.3772 | 0.3736 | 0.1866 | 0.9980 | 0.4973 | 0.4990 | 0.9580 |
| HLD97 | 0.6612 | 0.3388 | 0.6026 | 0.3974 | 0.3477 | 0.1285 | 0.8665 | 0.4480 | 0.4230 | 0.2579 |
| HLD99 | 0.1530 | 0.8470 | 0.4188 | 0.5812 | 0.2256 | 0.0501 | 0.6802 | 0.2592 | 0.2649 | 0.7830 |

As shown in Fig 2, the Heat map of allele frequencies was drawn by *R* statistical software v 3.0.2 on the basis of insertion allele frequencies of the 30 InDel loci to describe the relationships between 30 InDel loci (horizontal axis) and different populations (vertical axis). Red represented high insertion frequencies which meant relative low deletion frequencies. On the contrary, blue represented low insertion frequencies which meant relative high deletion frequencies. In the graph, HLD 118, HLD 99, HLD 64,HLD 81, HLD 67 and HLD 84 loci showed high insertion frequencies in East Asia populations which contained Han populations in three different regions (Neuvonen et al., 2012, Wang et al., 2014, Hong et al., 2013) Yi (Zhang et al., 2015), Xibe (Meng et al., 2015), South Korean (KM et al., 2014), Tibet Tibetan and Qinghai Tibetan [7], She (Wang et al., 2014)and Tujia groups (Shen et al., 2016), while HLD 111, HLD 39, HLD 114 and HLD 122 loci were shown low insertion frequencies in those groups. Besides, six Mexican (Martníez-Cortés et al., 2016) groups expressed the tendency to have high insertion frequencies in HLD 83, HLD 58, HLD 125 loci and low insertion frequencies in HLD 64 and HLD 81 loci. Furthermore, clustering analysis of 30 InDel loci was conducted as well, 30 InDel loci were respectively placed into two large clusters (I and II) shown in the Heat map, the cluster I contained 10 loci, while the cluster II included 20 loci.

**Fig 2.**
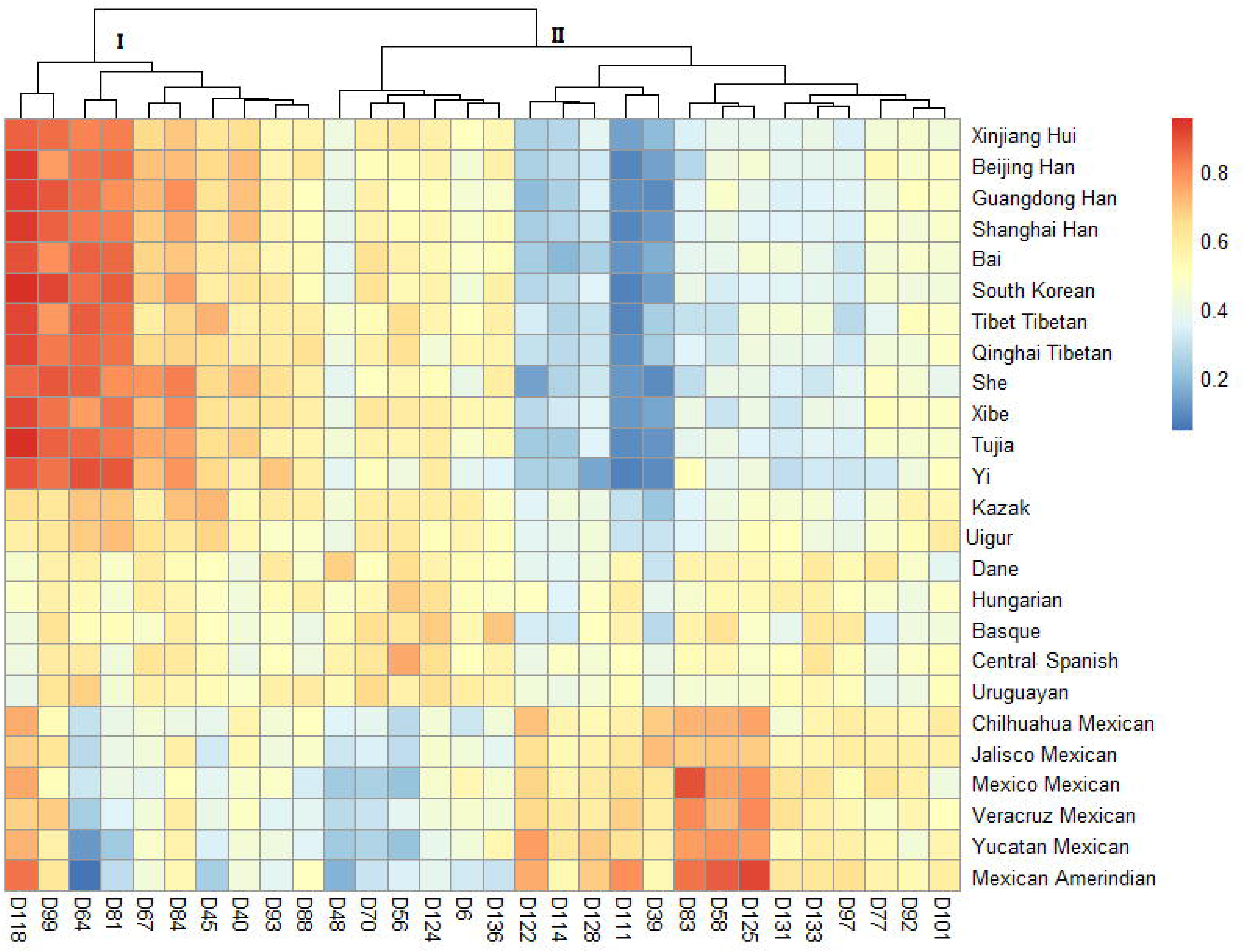
Heat map of allele frequency of Xinjiang Hui and 24 previously studied populations on the basis of R software

### 3.3 Population differentiations based on 30 InDels

We applied the analysis of molecular variance (AMOVA) method using the ARLEQUIN software version 3.1 to compare the allele differentiations between Xinjiang Hui group with other 24 previously reported populations based on *p* values (data not shown).The minimum numbers of significant differences observed at 30 InDel loci were found between Xinjiang Hui group and Bai at 2 loci; Qinghai Tibetan at 3 loci, respectively. On the contrary, the maximum numbers were observed between Xinjiang Hui and six Mexican groups, including Chihuahua Mexico, Jalisco Mexican, Mexican Amerindian, Yucatan Mexican, Veracruz Mexican, and Mexico Mexican at 27, 27, 27, 27, 26, and 26 loci, respectively. Besides, HLD 39, HLD 40, HLD 99, HLD 111, HLD 118, HLD 84 loci showed relatively high ethnic diversities, with the extraordinary distinction between Xinjiang Hui group and the other 17, 17, 17, 18, 18, and 19 populations. Whereas, HLD 92, HLD 6, HLD 101, HLD 88 and HLD 136 loci showed relatively low ethnic diversities with the significant differentiation among Xinjiang Hui group and other 4, 6, 7, 7 and 7 populations.

As shown in Fig 3, the Heat map was generated by *Fst* values of pairwise population differentiations, which reflected population genetic relationships by the depth of every block’s color. We used the R statistical software to draw a Heat map of *Fst* values between Xinjiang Hui and other 24 population. The genetic divergence among populations was represented by a shade of color green. The deeper color represented the bigger *Fst* value, and meant the more, genetic differentiation, whereas the lighter color indicated the lower *Fst* values and the less genetic differentiation.

**Fig 3.**
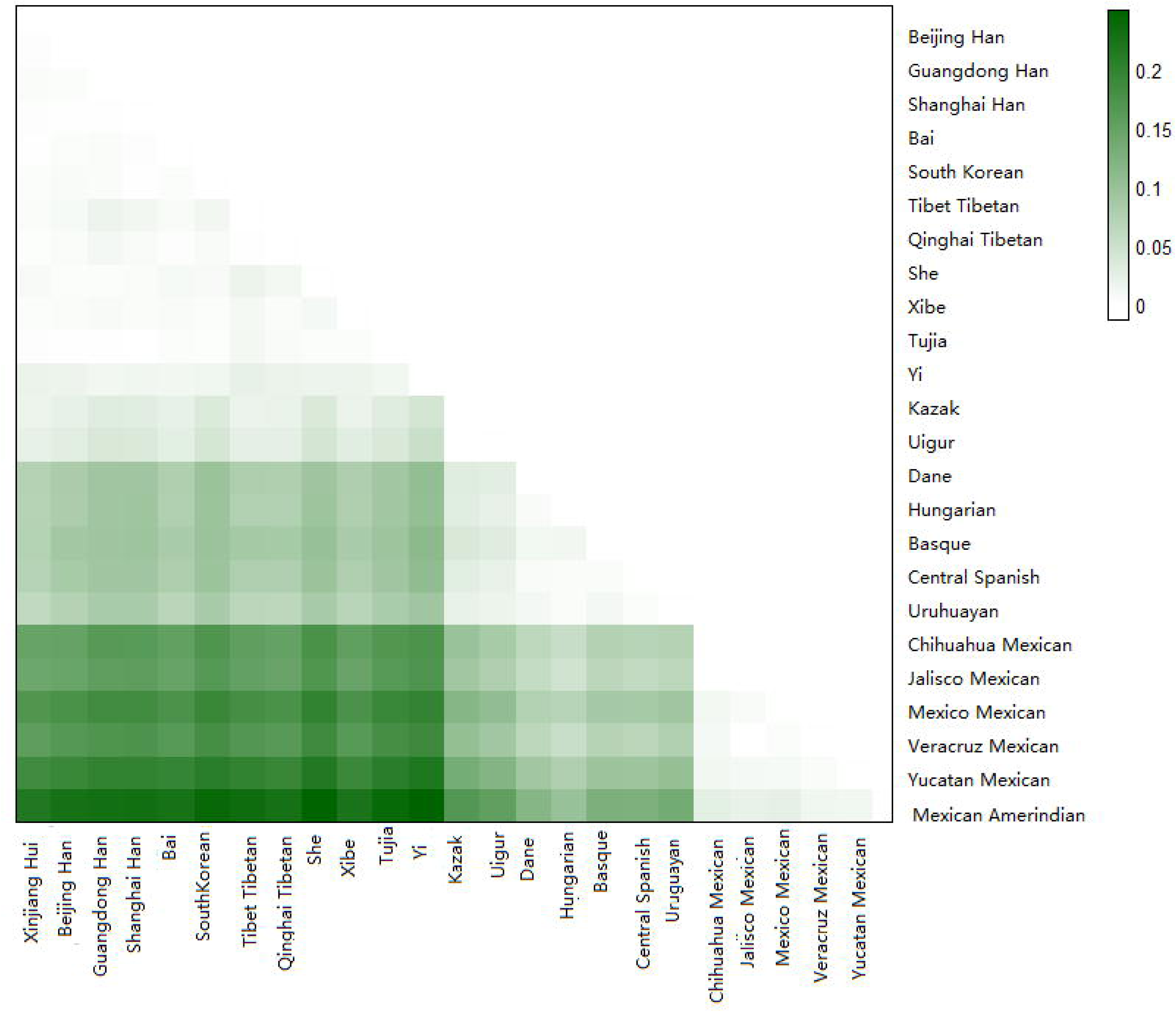
Heat map of pairwise of *Fst* of Xinjiang Hui and 24 previously studied populations on the basis of R software

We calculated the genetic distance based on the same set of 30 InDel loci by inspecting genetic divergence of Xinjiang Hui and other 24 populations and use the parameter *D*_*A*_ to presented it. The *D*_*A*_ values were shown in S1Table, the minimum *D*_*A*_ values was 0.0011 (Xinjiang Hui and Bai, Xinjiang Hui and Shanghai Han), 0.0012 (Xinjiang Hui and Qinghai Tibetan); and the diminutive genetic distances were distinguished between the Xinjiang Hui group and most of the East Asia groups such as Xibe and Tujia (0.0013), Beijing Han (0.0015), South Korean (0.0017), Guangdong Han (0.0018), Tibet Tibetan (0.0021), She (0.0027). Nevertheless, the further genetic distances were discovered between the Xinjiang Hui and Yi (0.0055), Kazak (0.006), Uigur (0.0072) populations, besides the grander genetic distances were found among Uruguayan (Saiz et al., 2014) (0.0174), Dane (SL et al., 2012) (0.0202), Central Spanish (Martín et al., 2013) (0.0199), Basque [18] (0.0205), and Hungarian (Z et al., 2012) (0.0201) populations, while the maximum *D*_*A*_ values (>0.397) were found with six Mexican groups.

### 3.4 Phylogenetic reconstruction of Xinjiang Hui and other populations

In order to reveal the population genetic relationships among the 25 populations, two different methods were selected to conduct the phylogenetic reconstruction using MAGA software v 5.0 and PHYLIP software v 3.6, respectively. An unrooted tree conducted by PHYLIP software v 3.6 was shown in Fig 4, which revealed the relationships between Xinjiang Hui group and the 24 previously published populations. All the 25 populations were divided into three major branches, the top side was East Asian populations including South Korean and almost Chinese populations such as three Han populations, Tibetan groups in two different regions, Xinjiang Hui, Bai, Xibe, Yi, Tujia, She groups; the lower side included six Mexican groups, Uruguayan, Basque, Central Spanish, Dane and Hungarian populations; and Kazak and Uigur groups were found in the middle, between these two branches. Xinjiang Hui group trended to be much closer to East Asian populations. Another phylogenetic tree conducted by MAGA software v 5.0 was reconstructed based on the *D*_*A*_ distances among the 25 populations. As shown in Fig 5, there were two main branches, the first one included the six Mexican groups, and the second one contained the East Asian groups, Uigur, Kazak and European groups. In addition, we found that Xinjiang Hui first clustered with two Tibetan groups, and then clustered with other Chinese populations. In short, two phylogenetic dendrograms had similar result.

**Fig 4.**
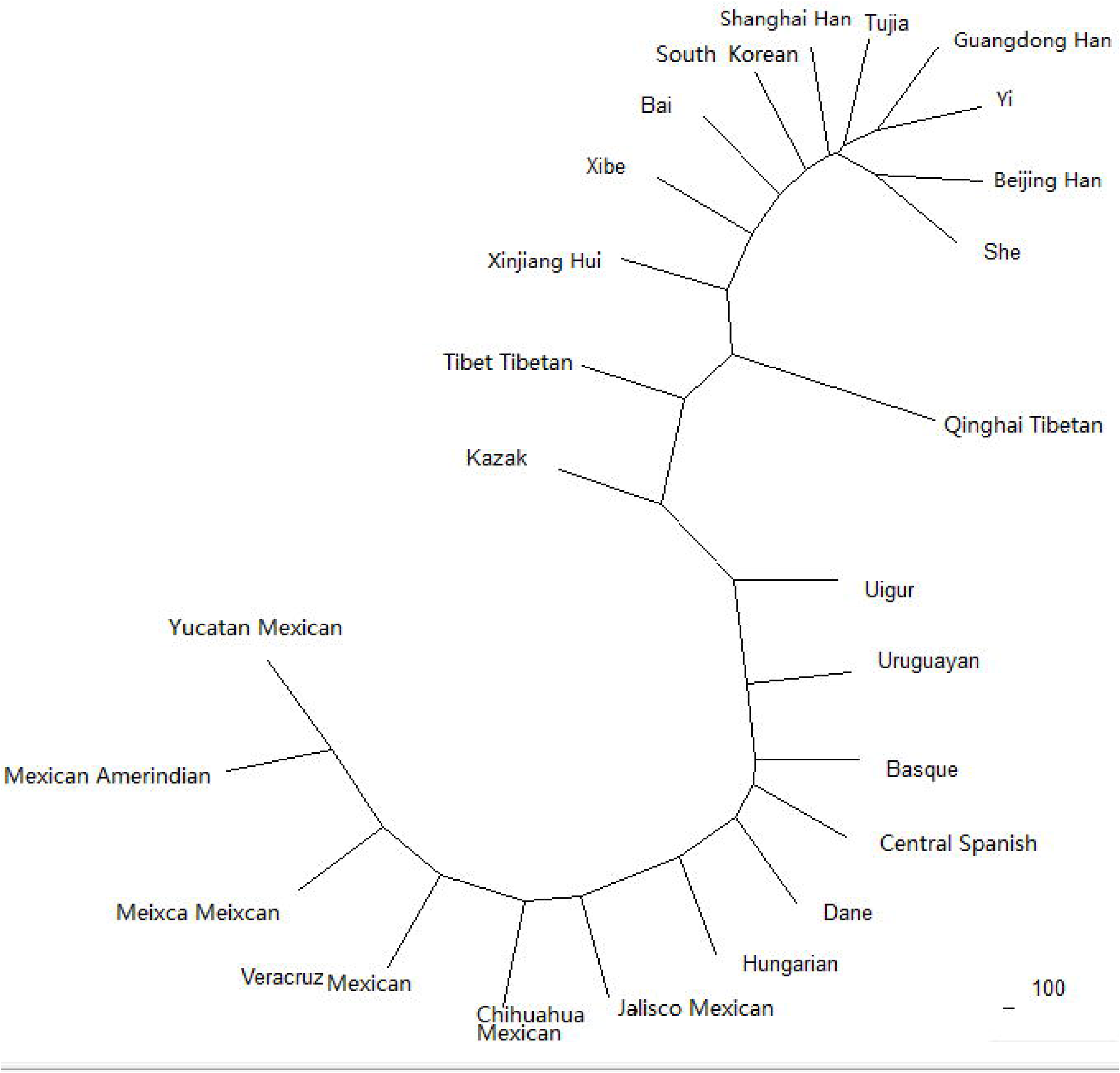
Phylogenetic trees generated to do the Analysis of phylogenetic relationships between Xinjiang Hui group and 24 previously studied populations on the basis of PHYLP software (version 3.6) based on allele frequency

**Fig 5.**
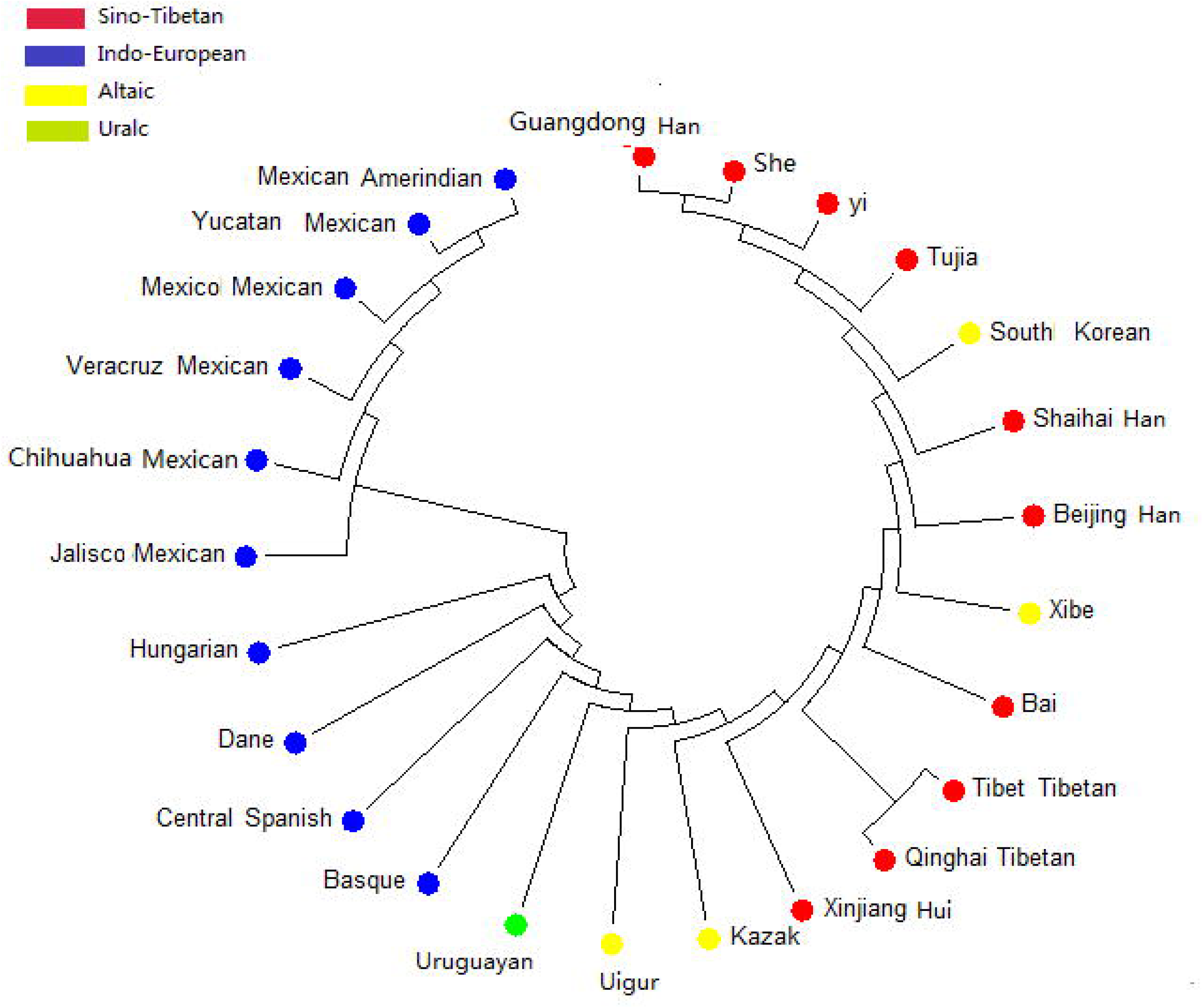
Phylogenetic trees generated to do the Analysis of phylogenetic relationships between Xinjiang Hui group and 24 previously studied populations on the basis of MAGA software (version5.0) based on DA values

### 3.5 Multi-Dimensional Scaling and Principal component analysis of Xinjiang Hui and other populations

In this study, the MDS plot was conducted based on the *Fst* values using SPSS18.0 software was conducted to characterize the genetic relationships between Chinese Xinjiang Hui group and the studied 24 populations. As shown in Fig 6, each population was represented by a small circle in the multidimensional space, and the distances between the small circles showed the genetic differentiation among the populations. There were four main population clusters in accordance with their geographic distributions divided clearly, including South Korean and most Chinese group; Uigur and Kazak group; six Mexican groups and European groups, respectively. The target population (Xinjiang Hui group) scattered in East Asian region, which mean that Xinjiang Hui group was more likely to have closer relationship with East Asian populations. In this study, PCA plot was carried out by the software of EIGENSOFT 6.0.1(AL et al., 2006), which could be shown in Fig 7. According to the geographic region, the whole 25 populations were partitioned into four regions: one dot represented a sample, and different colors meant different categories of populations. And the East Asian groups except Hui group, Uigur and Kazak, six Mexican, and European groups corresponded to the colors blue, pink, orange and green, respectively. And the Xinjiang Hui group was represented by black dots, a black dot instead of one sample with the sample number on it, and all the Xinjiang Hui individuals were located in East Asian area overwhelmingly, which suggested that Xinjiang Hui and East Asian groups had closer relationship.

**Fig 6.**
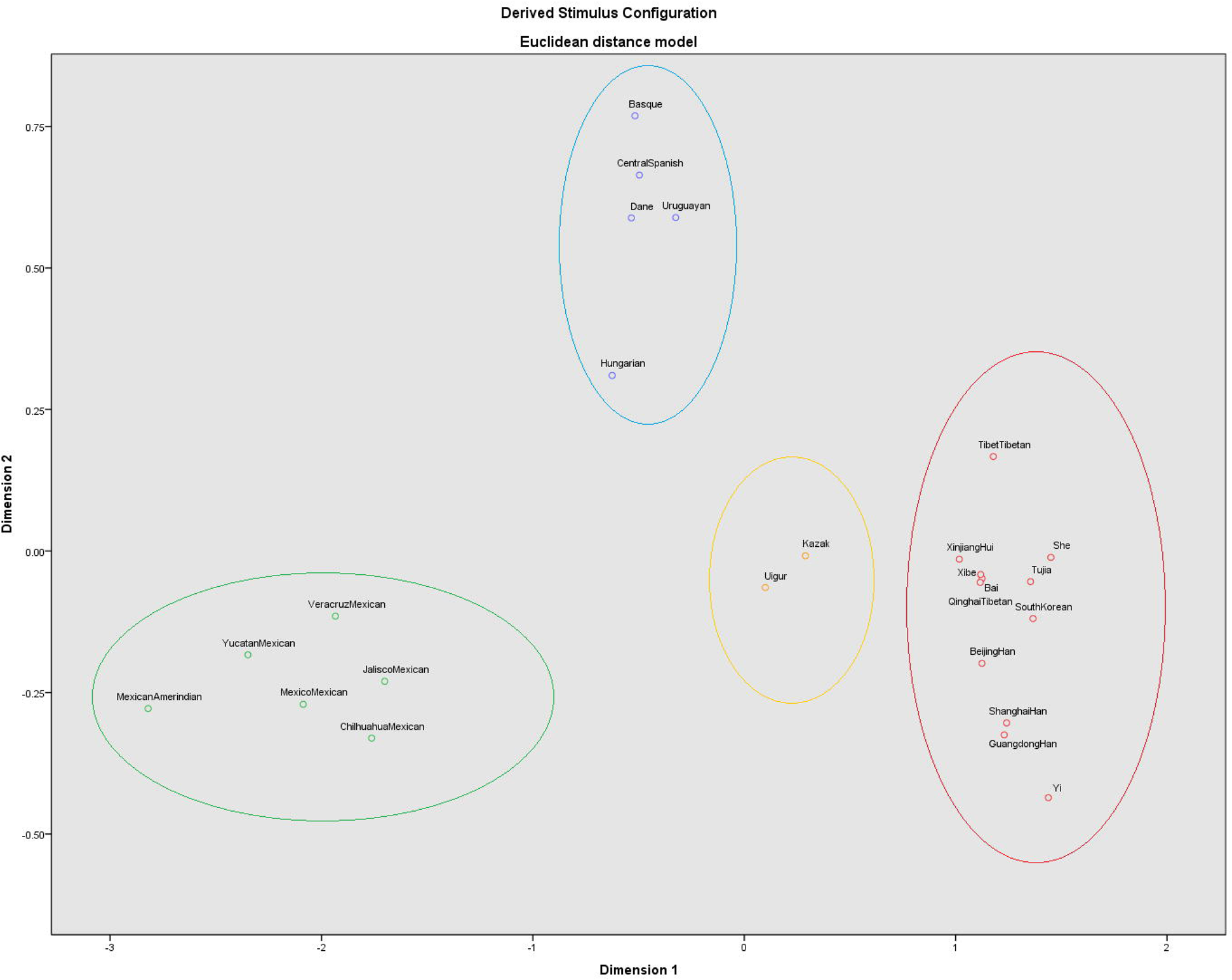
Multi-Dimensional Scaling of Xinjiang Hui group conducted by SPSS18.0

**Fig 7.**
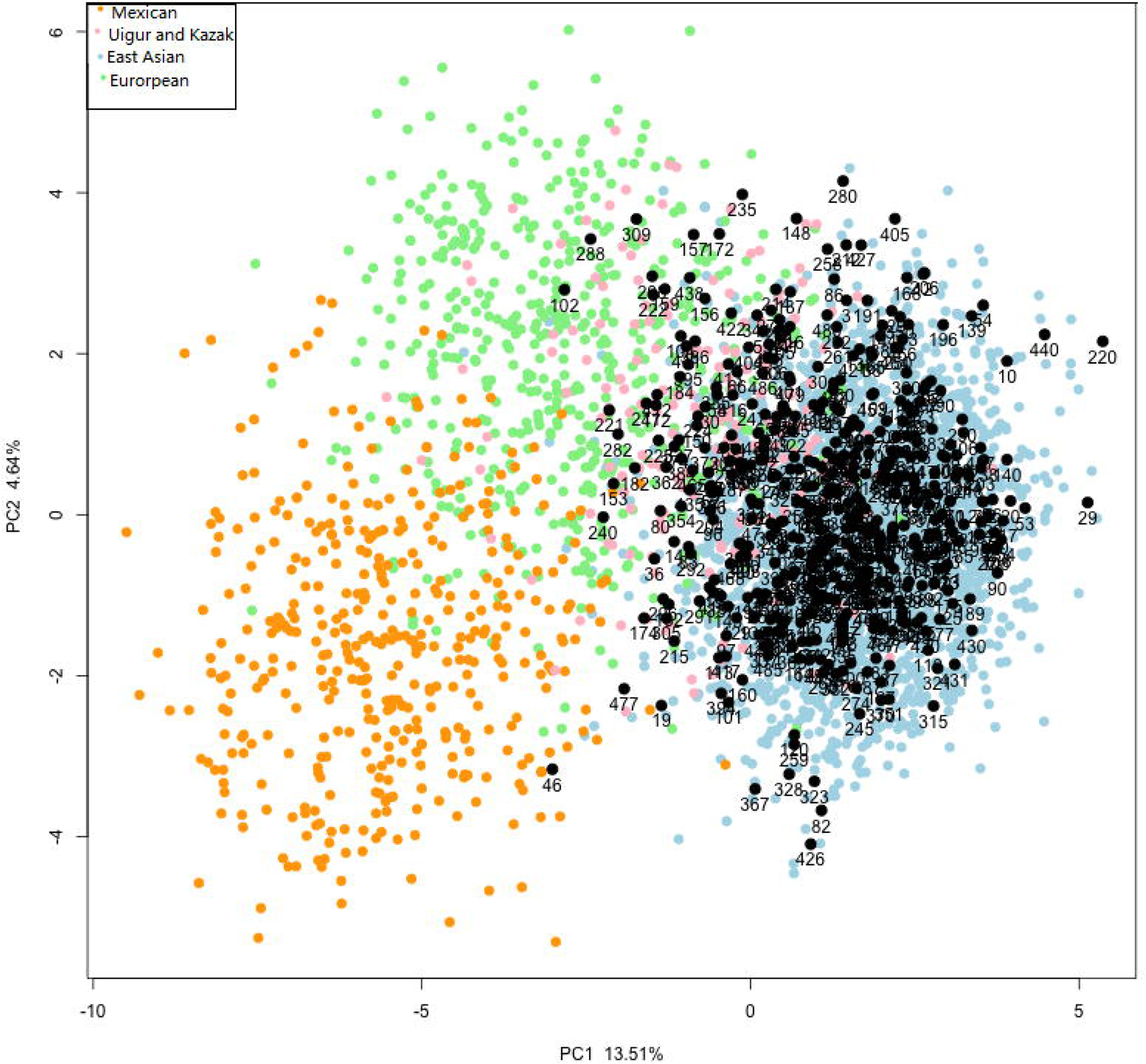
Principal component analysis with the method of software of EIGENSOFT 6.0.1 and MATLAB 2007a

### 3.6 Population structure analysis between Xinjiang Hui and other populations

We used STRUCTURE program v2.3.4 to appraise the genetic structure of Xinjiang Hui and the referenced 24 populations. As shown in Fig 8, the two vertical lines of black part represented the structure diagram of a population(Porrashurtado et al., 2013), when K=2, the most Asian groups were completely separated with European and Mexican groups, and the Asian groups were filled with the mixture red component while European and Mexican groups were filled with green. These ethnic groups had no definite subdivided into different clusters when K=3, the fourteen Asian groups; five European groups; and six Mexican groups can be distinguished clearly, which filled with mixture of green, yellow and blue; red and blue; and red, respectively. In addition,when K=6, Kazak and Uigur groups can be separated from populations Asian and European.

**Fig 8.**
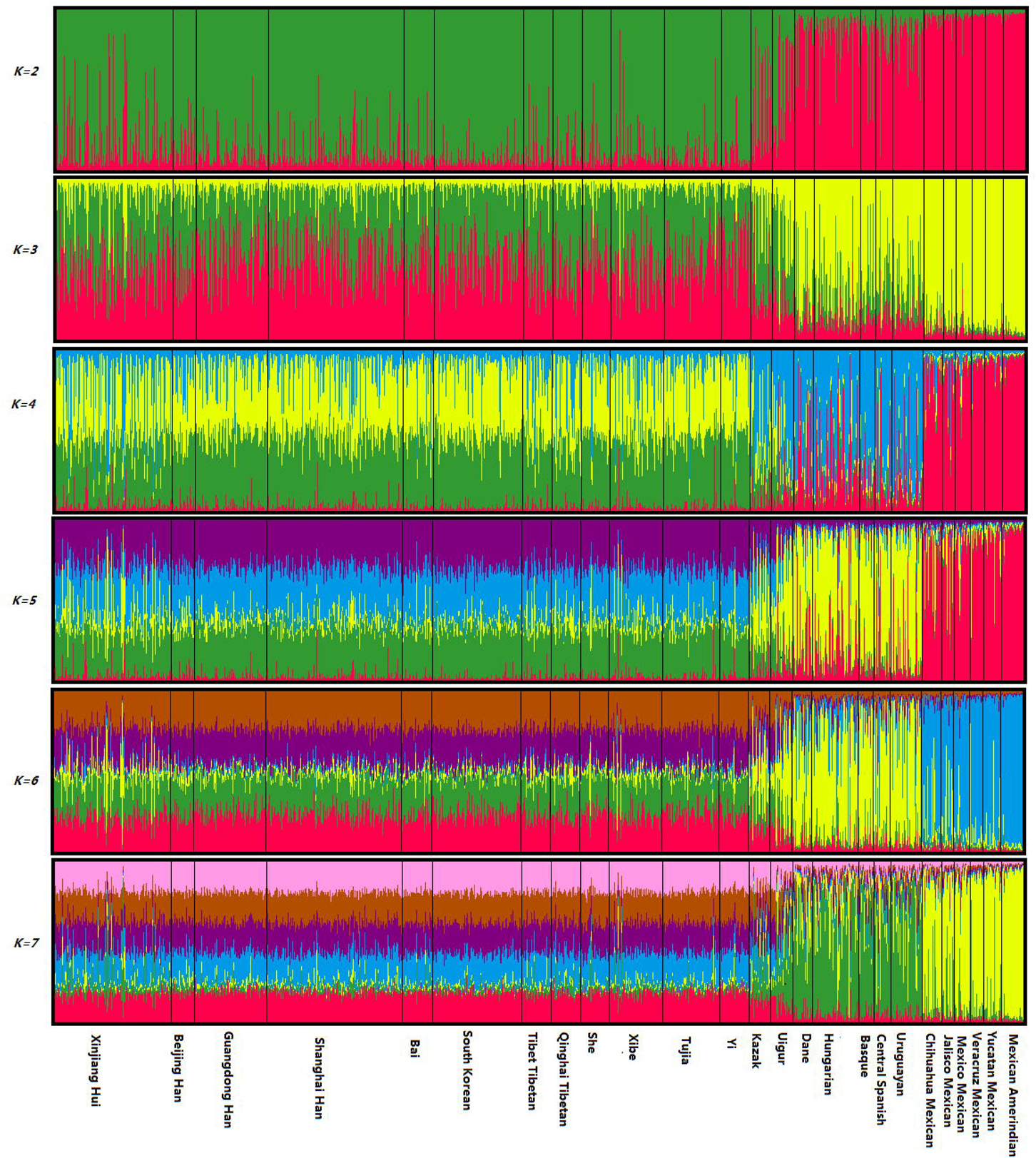
Structure analysis of Xinjiang Hui group based on STRUCTURE program v2.2.

## 4 Discussion

### 4.1 Linkage disequilibrium analysis

Before these 30 InDel loci were applied for forensic research, it was necessary to probe into linkage disequilibrium analysis.Previous study (Slatkin, 2008)clarified that in the same or two random chromosomes, any of the two loci showed no relevance, they were independent loci and could be qualified for population genetic analysis. In this study, there was no significant LD among the loci, which indicated independent relationship with each other of the 30 InDels in the Chinese Xinjiang Hui group.

### 4.2 Allelic polymorphisms and forensic statistical parameter analysis

A Heat map of allelic frequencies demonstrates data in the form of a rectangular matrix, and the color of each rectangle represents the value of the correspondent item in the matrix (Wilkinson and Friendly, 2009). In East Asians, 6 loci showed high insertion frequencies and 4 loci showed low insertion frequencies, and Xinjiang Hui group showed coincident allele frequencies at those loci. Besides, European and six Mexican populations also put up visible differences in 30 InDel loci. In short, the allele frequencies of 30 InDel loci could distinguish different populations to a certain degree, and the results indicated that Xinjiang Hui group was similar distritions of alleleic frequencies with the East Asians, especially most Chinese populations.

Generally speaking, in the forensic application field, CPE and CDP are common indicators to estimate the forensic practical efficiency of the 30 InDel loci. In the study, the CPE and CDP were 0.988849 and 0.99999999999378, respectively. The relatively lower CPE value indicated that the 30 loci could be used as complementary genetic markers for paternity testing.However, the high CDP value clarified that the 30 InDel loci could be regarded as efficient genetic markers in forensic identification cases.

### 4.3 Population differentiations based on 30 InDel loci

The *p* values of pairwise populations were conducted by the method of AMOVA to reveal genetic differentiations among the 25 populations, and the critical value of *p* is 0.05. When a *p* value is less than 0.05 (Bhattacharya and Habtzghi, 2002), it represents significant genetic differences between the two populations at the same locus. As for the population, we observed that Xinjiang Hui group had least discrepancies with Bai, Qinghai Tibetan and Shanghai Han, and so on; on the contrary, the maximum numbers of significant differences observed at 30 loci were found between Xinjiang Hui and six Mexican groups, which meant those groups had significant genetic differences and farthest genetic distances with Xinjiang Hui group. Besides, we made a further analysis in different loci, and HLD39, HLD40, HLD99, HLD111, HLD118, HLD84 loci showed relatively high ethnic diversities, while HLD92, HLD6, HLD101, HLD88, HLD136 loci showed relatively low ethnic diversities. The above results indicated that allele frequencies data of 30 InDels loci had significant differences in different ethnic groups, so it was very necessary and had practical meaning for forensic application researches in different populations. In addition, on the basis of the 30 InDel loci, we carried out a Heat map based on *Fst* distances of pairwise populations to make further disclosure of the relationships among the 25 populations. By analyzing the Heat map, we observed that the 25 populations could be divided into three classes mainly of relying on the depth of color: the East Asians, Europeans and Mexicans. We could infer that South Korean and the most Chinese groups such as Bai, three Han populations, Tujia, two Tibetan groups and Xibe had the relative lighter color for their pairwise *Fst* values, and which mean the above groups had closer genetic relationships with Xinjiang Hui group, and vice versa.

Genetic distance is considered as an effective analysis method to reveal the genetic divergence between populations within a species (Slatkin, 2008). It’s also a useful strategy to study the origin and history of different populations. In general, populations with the similar allelic distributions mean that they have smaller genetic distances. In the study, small genetic distances were found between Xinjiang Hui and most Chinese populations, it meant that they might have genetic affinity or common ancestor to a certain extent, and the present results was coincident with the previous study based on 26 Y-STR loci, which Zhao et al.made the conclusion that Hui group showed litter differences with most Chinese groups (Zhao et al., 2017).

### 4.4 Phylogenetic analysis and PCA and MDS among 25 populations

The aim of phylogenetic analysis is to infer or evaluate the relationships between different populations. In the present study, two different phylogenic trees were reconstructed to reveal the genetic relationships among the 25 populations based on *D*_*A*_ values (conducted by MAGA software v 5.0) and allelic frequencies (PHYLIP software v 3.6) of 30 InDel loci, respectively. The results indicated that the Chinese Xinjiang Hui group was similar with Han populations, Tibetan, and other Chinese populations at the molecular level. The PCA and MDS figures also reflected the similar genetic relationships between Xinjiang Hui and 24 populations. PCA is a statistical method which is mostly used as a tool in exploratory data analysis and for making predictive models (Hotelling, 1933). MDS is a multivariate statistical data analysis method that reveals the similarities or differences of distance-like data as a geometrical picture, and the multidimensional scaling analysis can commendably expound the possibility space in increasing the sesparability of the clusters (Borg and Groenen, 2003). According to the cluster analysis of two trees,Xinjiang Hui group was more likely to have close relationships with most Chinese groups and South Korean, especially Han groups and Tibetan groups. This result also gained the support of historical record and archaeological study. According to Jaschok et al (CHIARA, 2002) during Tang Dynasty, many Muslims from Persia, Arab and Central Asian entered China through Xinjiang via Silk Road or Maritime Silk Route. From then on, the immigrations came from Arab, Persia and Central Asian due to business or warfare. Then the Hui group continued to interact with the Mongolian, the Uyghur and the Han groups. In addition, from Yuan dynasty to Qing dynasty, there are many Hans and few northwest ethnic groups joined the Hui group because of marriage and reclamation. In the long tern of mixed dwelling with the Hans, and particularly due to the augment number of Hans joining their group, Hui abandoned their original languages such as Arab and Persian, and tendency to speak the Han language only, moreover, Hui group gradually assimilate the Han’s culture. The above results also confirmed that the Hui group and Han population have more close relationships. (Lipman, 1998)

### 4.5 Population structure analysis among 25 populations

The population structure analysis was aimed at describing the clusters of different populations based on their raw genotypes at a set number of loci using a Bayesian algorithm, and it could make the identification of genetically homogeneous populations of individuals (Porrashurtado et al., 2013). When populations were far from each other in geographical distance or consanguinity such as Asian and American, the individuals in such populations generally had various memberships coefficient in deductive cluster (Earl and Vonholdt, 2012). According to the specification of structure program, similar structure atlas always indicated close memberships. Compared with 10 East Asian populations, Kazak and Uigur, 5 European populations and 6 Mexican populations, the Chinese Xinjiang Hui were more similar with the East Asians, which had the similar results with the MDS, PCA and the phylogenetic trees. Therefore, based on structure analysis by raw data of 30 Indel loci, the Chinese Xinjiang Hui group was homogeneous with the previous studied East Asian groups.

## 6 Conclusion

In this study, we evaluated the allele frequencies and forensic parameters of the autosomal 30 InDels as new tremendous potential diallelic genetic markers in population genetic studies and forensic sciences. According to the above results, Xinjiang Hui group had close genetic relationships with most Chinese populations especially Han populations based on the results of population differentiations, principal component analysis, population structure analysis, and phylogenetic analysis. Aiming to further reveal the genetic background and origin of Xinjiang Hui group, further study, such as autosomal ancestry-informative InDel markers should be performed in our later research.

## Acknowledgments

We thank Drs Pingming Qiu (Southern Medical University), Bofeng Zhu (Southern Medical University) for doing research design, and collecting a large number of samples. Tong Xie (Southern Medical University) and Yuxin Guo (Xi’an Jiaotong University) kindly performed the forensic data analysis. Ling Chen (Southern Medical University), Yunchun Tai(Southern Medical University), and Yating Fang (Southern Medical University) kindly performed the analysis of contrast populations. We sincerely thank Southern Medical University for providing the research platform.

## Author contributions

T.X, Y.G, P.Q and B.Z. wrote the main manuscript text, Y.T, Y.Z, Y.F, and L.C, did the data processing and the manuscript modification, T.X, and Y.G prepared the figures. All authors reviewed the manuscript.

## Supporting information

S1 Fig The LD analysis schema between the 30 InDel loci using the SNPAnalyzer 2.0 program

S1 Table The values of DA of pairwise populations among Chinese Xinjiang Hui group and referenced populations

